## Supplementary Materials for "The effects of 17α-estradiol treatment on endocrine system revealed by single-nucleus transcriptomic sequencing of hypothalamus"

**Figure legends**

**Figure 2—figure supplement 1.** Top 20 signaling pathways or gene sets significantly positively or negatively associated with MitoCarta OXPHOS subunits in Neuron.O. Significant *p*-values (<0.05) are indicated by a star.

**Figure 2—figure supplement 2.** **The variable response patterns of non-neuron cells to aging and** **17α-estradiol treatment in hypothalamus.** Dot plot of overall expression levels of selected pathways from the two opposing signaling networks in 9 non-neural cell types.

**Figure 2—figure supplement 3. The top enriched pathways of significantly expressed genes in Micro, Astro and Neuron between O.T and O.** Top 12 enriched GOBP pathways via DAVID Functional Annotation Tools in Micro (O vs Y, O.T vs O), Astro (O vs Y, O.T vs O), and Neuron (O vs Y, O.T vs O) in significantly down-regulated or up-regulated genes, which were calculated via FindMarker function in R package Seurat (test.use = bimod, min.pct = 0.1, logfc.Threshold = 0.25).

**Figure 3—figure supplement 1.** **The similarity of neuropeptide-expressing subclusters or receptor-expressing subclusters in young rat hypothalamus.** **(A, B)** Heatmaps showing the similarity of neuropeptide-expressing subclusters (A) and receptor-expressing subclusters (B) in the hypothalamus of young rats (left panels). Each subcluster contains no fewer than 10 cells. Venn diagrams (right panels) display the overlap of cell barcodes among neuronal subclusters with higher similarity.

**Figure 6—figure supplement 1. The expression profiles of selected pathways from the two opposing signaling networks in 6 cardiovascular system-related neurons.** Among the 26 selected pathways, 8 were metabolism-related pathways and usually elevated during aging (orange) and 18 were either negatively correlated with MitoCarta OXPHOS subunits or pathways related to synapse activity (dark green).

**Figure 6—figure supplement 2. Bidirectional Two-sample MR analysis of causal effects between 203 endocrine-related factors and Oxt (id: prot-a-2159).** **(A)**	Significant causal effects (p <0.05, IVW) related to exposure Oxt (id: prot-a-2159) and 203 endocrine-related outcomes in both bidirectional MR analysis. **(B)**	Significant causal effects (p <0.05, IVW) related to 203 endocrine-related exposures and outcome Oxt (id: prot-a-2159) in both bidirectional MR analysis.

**Figure 7—figure supplement 1.** The top 20 and bottom 20 neuron subtypes based on the mean expression values of c4-up-signature, ranked by the values in sample O.T, in neuropeptide- or hormone-secreting subtypes (upper panel) and in neuron subtypes expressing neuropeptide receptors or hormone receptors (lower panel).

**Figure 7—figure supplement 2. Two-sample MR analysis of causal effects of 203 endocrine-related exposures on outcome GNRH1. (A)** Significant causal effects (p <0.05, IVW), which were not significant in reverse MR analysis between 203 endocrine-related exposures and outcome GNRH1 (id: prot-a-1233). **(B)** Significant causal effects (p <0.05, IVW) in both directions of MR analysis.
