## Supplemental Data 6 for "The effects of 17α-estradiol treatment on endocrine system revealed by single-nucleus transcriptomic sequencing of hypothalamus"

**A**

| outcome | method | nsnp | OR(95%CI) | pval | Q_pval | pleiotropy.pval |
|---|---|---|---|---|---|---|
| Orexigenic neuropeptide QRFP \|\| id:prot−a−2467 | IVW | 68 | 1.352 (1.286 to 1.422) | 1.3e−31 | 1 | 0.26 |
| Insulin growth factor−like family member 3 \|\| id:prot−a−1453 | IVW | 34 | 1.578 (1.417 to 1.756) | 8.9e−17 | 0.000046 | 0.21 |
| Insulin−like growth factor I \|\| id:prot−a−1443 | IVW | 34 | 1.313 (1.222 to 1.410) | 8.1e−14 | 0.88 | 0.4 |
| Agouti−related protein \|\| id:prot−a−59 | IVW | 68 | 1.182 (1.124 to 1.244) | 8.5e−11 | 0.58 | 0.38 |
| Tachykinin−4 \|\| id:prot−a−2922 | IVW | 34 | 1.270 (1.177 to 1.371) | 8.6e−10 | 0.26 | 0.85 |
| Amyloid−like protein 2 \|\| id:prot−a−122 | IVW | 34 | 1.246 (1.160 to 1.338) | 1.6e−09 | 0.77 | 0.38 |
| Leptin receptor \|\| id:prot−a−1725 | IVW | 34 | 1.252 (1.164 to 1.348) | 2e−09 | 0.38 | 0.61 |
| Serpin I2 \|\| id:prot−a−2702 | IVW | 34 | 1.323 (1.204 to 1.453) | 6e−09 | 0.0053 | 0.26 |
| Gastrin−releasing peptide \|\| id:prot−a−1278 | IVW | 34 | 1.233 (1.148 to 1.324) | 9.4e−09 | 0.68 | 0.12 |
| Amyloid beta A4 protein \|\| id:prot−a−139 | IVW | 34 | 1.231 (1.146 to 1.322) | 1.2e−08 | 0.92 | 0.5 |
| Melanocortin−2 receptor accessory protein \|\| id:prot−a−1936 | IVW | 34 | 1.307 (1.192 to 1.433) | 1.3e−08 | 0.0095 | 0.91 |
| Estradiol 17−beta−dehydrogenase 2 \|\| id:prot−a−1383 | IVW | 34 | 1.281 (1.175 to 1.396) | 1.8e−08 | 0.045 | 0.5 |
| Clusterin \|\| id:prot−a−596 | IVW | 34 | 1.267 (1.156 to 1.388) | 4e−07 | 0.012 | 0.21 |
| Brain−derived neurotrophic factor \|\| id:prot−a−242 | IVW | 34 | 1.198 (1.116 to 1.287) | 7e−07 | 0.59 | 0.18 |
| Proprotein convertase subtilisin/kexin type 7 \|\| id:prot−a−2212 | IVW | 34 | 1.193 (1.111 to 1.281) | 1.3e−06 | 0.66 | 0.22 |
| Insulin−like growth factor−binding protein−like 1 \|\| id:prot−a−1452 | IVW | 34 | 1.175 (1.087 to 1.270) | 0.000049 | 0.22 | 0.77 |
| Melanocortin−2 receptor accessory protein 2 \|\| id:prot−a−1935 | IVW | 34 | 1.144 (1.065 to 1.229) | 0.00025 | 0.45 | 0.44 |
| Prolactin receptor \|\| id:prot−a−2377 | IVW | 34 | 1.236 (1.103 to 1.386) | 0.00027 | 2.3e−06 | 0.33 |
| Atrial natriuretic factor \|\| id:prot−a−2076 | IVW | 34 | 1.153 (1.066 to 1.247) | 0.00038 | 0.2 | 0.32 |
| Pituitary adenylate cyclase−activating polypeptide \|\| id:prot−a−42 | IVW | 34 | 1.148 (1.061 to 1.242) | 0.00062 | 0.18 | 0.36 |
| Tachykinin−3 \|\| id:prot−a−2921 | IVW | 34 | 1.126 (1.048 to 1.210) | 0.0011 | 0.72 | 0.26 |
| Corticotropin \|\| id:prot−a−2326 | IVW | 34 | 1.127 (1.039 to 1.222) | 0.0038 | 0.12 | 0.67 |
| Insulin−like peptide INSL6 \|\| id:prot−a−1563 | IVW | 34 | 1.124 (1.030 to 1.226) | 0.0084 | 0.038 | 0.34 |
| Prolactin \|\| id:prot−a−2375 | IVW | 34 | 1.097 (1.021 to 1.178) | 0.011 | 0.49 | 0.39 |
| Serpin A12 \|\| id:prot−a−2690 | IVW | 34 | 1.146 (1.003 to 1.309) | 0.046 | 5.3e−11 | 0.97 |

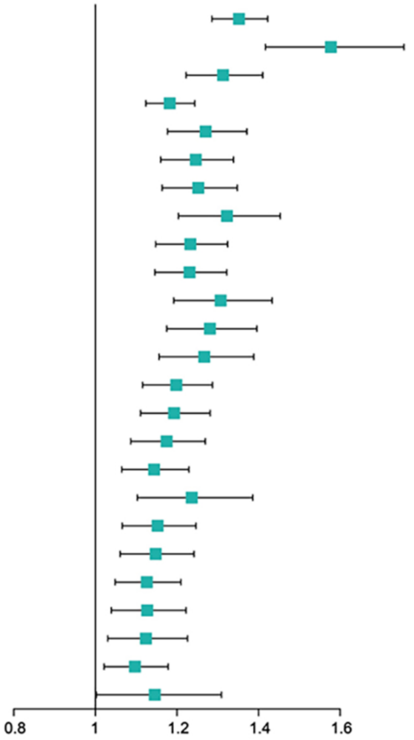


**B**

| exposure | method | nsnp | OR(95%CI) | pval | Q_pval | pleiotropy.pval |
|---|---|---|---|---|---|---|
| Insulin growth factor−like family member 3 \|\| prot−a−1453 | IVW | 37 | 1.413 (1.325 to 1.507) | 8.2e−26 | 0.54 | 0.6 |
| Estradiol 17−beta−dehydrogenase 2 \|\| prot−a−1383 | IVW | 33 | 1.325 (1.224 to 1.434) | 3.4e−12 | 0.24 | 0.7 |
| Insulin−like growth factor I \|\| prot−a−1443 | IVW | 32 | 1.298 (1.198 to 1.408) | 2.3e−10 | 0.15 | 0.81 |
| Orexigenic neuropeptide QRFP \|\| prot−a−2467 | IVW | 18 | 1.378 (1.246 to 1.525) | 5.1e−10 | 0.67 | 0.42 |
| Insulin−like peptide INSL6 \|\| prot−a−1563 | IVW | 22 | 1.363 (1.224 to 1.518) | 1.7e−08 | 0.085 | 0.84 |
| Gastrin−releasing peptide \|\| prot−a−1278 | IVW | 27 | 1.285 (1.173 to 1.409) | 8e−08 | 0.15 | 0.089 |
| Amyloid beta A4 protein \|\| prot−a−139 | IVW | 23 | 1.291 (1.176 to 1.418) | 8.1e−08 | 0.3 | 0.81 |
| Tachykinin−4 \|\| prot−a−2922 | IVW | 27 | 1.261 (1.147 to 1.386) | 1.6e−06 | 0.087 | 0.56 |
| Insulin−like growth factor−binding protein−like 1 \|\| prot−a−1452 | IVW | 18 | 1.277 (1.155 to 1.411) | 1.6e−06 | 0.63 | 0.95 |
| Brain−derived neurotrophic factor \|\| prot−a−242 | IVW | 25 | 1.226 (1.128 to 1.333) | 1.6e−06 | 0.89 | 0.21 |
| Tachykinin−3 \|\| prot−a−2921 | IVW | 27 | 1.210 (1.116 to 1.312) | 3.8e−06 | 0.56 | 0.51 |
| Proprotein convertase subtilisin/kexin type 7 \|\| prot−a−2212 | IVW | 25 | 1.191 (1.100 to 1.289) | 0.000016 | 0.86 | 0.59 |
| Prolactin \|\| prot−a−2375 | IVW | 33 | 1.183 (1.093 to 1.281) | 0.000033 | 0.27 | 0.37 |
| Amyloid−like protein 2 \|\| prot−a−122 | IVW | 17 | 1.216 (1.108 to 1.334) | 0.000035 | 0.97 | 0.22 |
| Clusterin \|\| prot−a−596 | IVW | 27 | 1.215 (1.106 to 1.334) | 5e−05 | 0.13 | 0.81 |
| Melanocortin−2 receptor accessory protein \|\| prot−a−1936 | IVW | 20 | 1.223 (1.093 to 1.369) | 0.00046 | 0.08 | 0.11 |
| Corticotropin \|\| prot−a−2326 | IVW | 23 | 1.194 (1.080 to 1.319) | 0.0005 | 0.16 | 0.41 |
| Atrial natriuretic factor \|\| prot−a−2076 | IVW | 29 | 1.150 (1.063 to 1.244) | 0.0005 | 0.83 | 0.62 |
| Pituitary adenylate cyclase−activating polypeptide \|\| prot−a−42 | IVW | 19 | 1.265 (1.108 to 1.445) | 0.00051 | 0.015 | 0.77 |
| Agouti−related protein \|\| prot−a−59 | IVW | 32 | 1.162 (1.067 to 1.265) | 0.00053 | 0.089 | 0.54 |
| Leptin receptor \|\| prot−a−1725 | IVW | 28 | 1.139 (1.048 to 1.239) | 0.0023 | 0.33 | 0.4 |
| Serpin I2 \|\| prot−a−2702 | IVW | 18 | 1.219 (1.072 to 1.387) | 0.0025 | 0.042 | 0.26 |
| Prolactin receptor \|\| prot−a−2377 | IVW | 15 | 1.241 (1.069 to 1.441) | 0.0046 | 0.0084 | 0.84 |
| Melanocortin−2 receptor accessory protein 2 \|\| prot−a−1935 | IVW | 31 | 1.092 (1.012 to 1.179) | 0.023 | 0.8 | 0.37 |
| Serpin A12 \|\| prot−a−2690 | IVW | 28 | 1.153 (1.005 to 1.322) | 0.042 | 3.8e−10 | 0.75 |

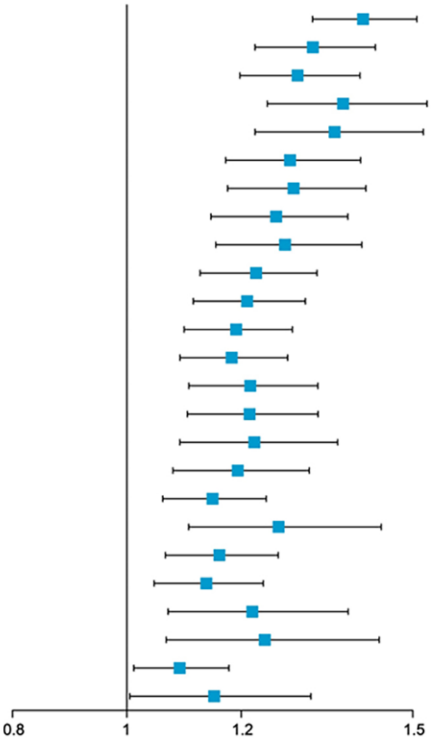
