## Supplemental Data 8 for "The effects of 17α-estradiol treatment on endocrine system revealed by single-nucleus transcriptomic sequencing of hypothalamus"

**A**

| exposure | outcome | method | nsnp | OR(95%CI) | pval | Q_pval | pleiotropy.pval |
|---|---|---|---|---|---|---|---|
| Thyrotropin−releasing hormone \|\| id:prot−a−3102 | GNRH1 | IVW | 30 | 1.163 (1.079 to 1.254) | 0.000085 | 0.73 | 0.51 |
| Agouti−related protein \|\| id:prot−a−59 | GNRH1 | IVW | 32 | 1.144 (1.064 to 1.231) | 0.0003 | 0.56 | 0.52 |
| Gastrin−releasing peptide \|\| id:prot−a−1278 | GNRH1 | IVW | 27 | 1.150 (1.061 to 1.247) | 0.00067 | 0.61 | 0.63 |
| Estradiol 17−beta−dehydrogenase 1 \|\| id:prot−a−1382 | GNRH1 | IVW | 21 | 1.183 (1.068 to 1.311) | 0.0013 | 0.24 | 0.36 |
| Tachykinin−3 \|\| id:prot−a−2921 | GNRH1 | IVW | 27 | 1.137 (1.039 to 1.244) | 0.0053 | 0.19 | 0.93 |
| Melanocortin−2 receptor accessory protein 2 \|\| id:prot−a−1935 | GNRH1 | IVW | 31 | 1.114 (1.032 to 1.202) | 0.0054 | 0.63 | 0.42 |
| Insulin−like growth factor−binding protein 2 \|\| id:prot−a−1448 | GNRH1 | IVW | 27 | 1.124 (1.035 to 1.221) | 0.0055 | 0.7 | 0.78 |
| Growth hormone levels \|\| id:ebi−a−GCST90010128 | GNRH1 | IVW | 13 | 1.064 (1.018 to 1.113) | 0.0064 | 0.74 | 0.43 |
| beta−nerve growth factor \|\| id:prot−a−2042 | GNRH1 | IVW | 17 | 1.166 (1.044 to 1.303) | 0.0065 | 0.28 | 0.67 |
| beta−nerve growth factor levels \|\| id:ebi−a−GCST90012042 | GNRH1 | IVW | 29 | 0.847 (0.744 to 0.964) | 0.012 | 0.37 | 0.92 |
| Cystatin−M \|\| id:prot−a−704 | GNRH1 | IVW | 27 | 1.104 (1.020 to 1.195) | 0.014 | 0.82 | 0.59 |
| Oxytocin−neurophysin 1 \|\| id:prot−a−2159 | GNRH1 | IVW | 34 | 1.091 (1.016 to 1.172) | 0.017 | 0.82 | 0.42 |
| Serpin I2 \|\| id:prot−a−2702 | GNRH1 | IVW | 18 | 1.145 (1.025 to 1.278) | 0.017 | 0.23 | 0.26 |
| Corticoliberin \|\| id:prot−a−658 | GNRH1 | IVW | 19 | 1.124 (1.021 to 1.238) | 0.017 | 0.87 | 0.96 |
| Pituitary adenylate cyclase−activating polypeptide \|\| id:prot−a−42 | GNRH1 | IVW | 19 | 1.125 (1.021 to 1.240) | 0.018 | 0.79 | 0.76 |
| Insulin−induced gene 1 protein \|\| id:prot−a−1559 | GNRH1 | IVW | 16 | 1.132 (1.019 to 1.257) | 0.021 | 0.58 | 0.99 |
| circulating leptin levels \|\| id:ebi−a−GCST90012076 | GNRH1 | IVW | 38 | 0.855 (0.740 to 0.987) | 0.032 | 0.88 | 0.42 |
| Neuromedin−B \|\| id:prot−a−2058 | GNRH1 | IVW | 23 | 1.094 (1.001 to 1.195) | 0.046 | 0.46 | 0.93 |
| growth hormone 1 \|\| id:prot−b−30 | GNRH1 | IVW | 5 | 0.865 (0.749 to 0.998) | 0.047 | 0.8 | 0.76 |
| Orexigenic neuropeptide QRFP \|\| id:prot−a−2467 | GNRH1 | IVW | 18 | 1.111 (1.000 to 1.234) | 0.049 | 0.37 | 0.54 |

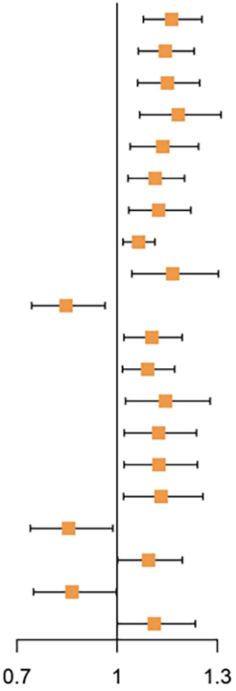


**B**

| exposure | outcome | method | nsnp | OR(95%CI) | pval | Q_pval | pleiotropy.pval |
|---|---|---|---|---|---|---|---|
| Glucagon \|\| id:prot−a−1181 | GNRH1 | IVW | 37 | 1.496 (1.372 to 1.630) | 5.3e−20 | 0.015 | 0.83 |
| Somatostatin−28 \|\| id:prot−a−2846 | GNRH1 | IVW | 27 | 1.760 (1.526 to 2.031) | 9.2e−15 | 4.3e−08 | 0.24 |
| Chromogranin−A \|\| id:prot−a−538 | GNRH1 | IVW | 24 | 1.659 (1.405 to 1.959) | 2.3e−09 | 1.7e−10 | 0.67 |
| Protachykinin−1 \|\| id:prot−a−2920 | GNRH1 | IVW | 32 | 1.478 (1.298 to 1.683) | 3.5e−09 | 9.4e−11 | 0.65 |
| Neuropeptide W \|\| id:prot−a−2082 | GNRH1 | IVW | 29 | 1.419 (1.246 to 1.617) | 1.4e−07 | 1.3e−11 | 0.068 |
| Corticotropin \|\| id:prot−a−2326 | GNRH1 | IVW | 23 | 1.206 (1.105 to 1.316) | 0.000027 | 0.9 | 0.53 |
| Secretogranin−1 \|\| id:prot−a−539 | GNRH1 | IVW | 18 | 1.212 (1.100 to 1.336) | 0.0001 | 0.64 | 0.16 |
| Atrial natriuretic factor \|\| id:prot−a−2076 | GNRH1 | IVW | 29 | 1.157 (1.069 to 1.252) | 0.00028 | 0.98 | 0.68 |
| Insulin−like growth factor I \|\| id:prot−a−1443 | GNRH1 | IVW | 32 | 0.848 (0.774 to 0.929) | 0.0004 | 0.017 | 0.47 |
| Estradiol 17−beta−dehydrogenase 2 \|\| id:prot−a−1383 | GNRH1 | IVW | 33 | 1.141 (1.061 to 1.228) | 0.00041 | 0.73 | 0.54 |
| Prolactin−releasing peptide \|\| id:prot−a−2376 | GNRH1 | IVW | 38 | 1.132 (1.033 to 1.239) | 0.0078 | 0.003 | 0.08 |
| Insulin−like growth factor 1 receptor \|\| id:prot−a−1444 | GNRH1 | IVW | 19 | 1.132 (1.031 to 1.243) | 0.0095 | 0.9 | 0.86 |
| Brain natriuretic peptide 32 \|\| id:prot−a−2077 | GNRH1 | IVW | 25 | 1.123 (1.028 to 1.227) | 0.01 | 0.35 | 0.84 |
| circulating leptin levels adjusted for BMI \|\| id:ebi−a−GCST90007324 | GNRH1 | IVW | 8 | 1.412 (1.036 to 1.926) | 0.029 | 0.85 | 0.55 |
| circulating leptin levels adjusted for BMI \|\| id:ebi−a−GCST90007322 | GNRH1 | IVW | 8 | 1.526 (1.032 to 2.256) | 0.034 | 0.89 | 0.67 |
| Progonadoliberin−2 \|\| id:prot−a−1234 | GNRH1 | IVW | 25 | 1.099 (1.002 to 1.207) | 0.046 | 0.18 | 0.56 |

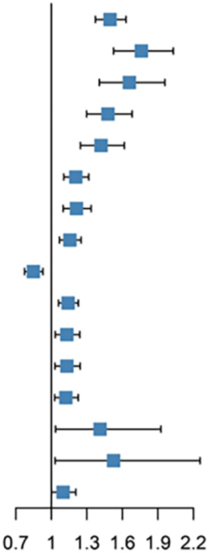
